# Genealogical distances under low levels of selection

**DOI:** 10.1101/495770

**Authors:** Elisabeth Huss, Peter Pfaffelhuber

## Abstract

For a panmictic population of constant size evolving under neutrality, Kingman’s coalescent describes the genealogy of a population sample in equilibrium. However, for genealogical trees under selection, not even expectations for most basic quantities like height and length of the resulting random tree are known. Here, we give an analytic expression for the distribution of the total tree length of a sample of size *n* under low levels of selection in a two-alleles model. We can prove that trees are shorter than under neutrality under genic selection and if the beneficial mutant has dominance *h* < 1/2, but longer for *h* > 1/2. The difference from neutrality is 𝒪 (*α*^2^) for genic selection with selection intensity *α* and 𝒪 (*α*) for other modes of dominance.

## 1 Introduction

Understanding population genetic models, e.g. the Wright-Fisher or the Moran model, can be achieved in various ways. Classically, allelic frequencies are described by diffusions in the large population limit, and for simple models such as two-alleles models, the theory of one-dimensional diffusions leads to predictions for virtually all quantities of interest (Ewens, 2004). Moreover, starting with Kingman (1982) and Hudson (1983), genealogical trees started to play a big role in the understanding of the models as well as of DNA data from a population sample. Most importantly, all variation seen in data can be mapped onto a genealogical tree. Under neutral evolution, the mutational process is independent of the genealogical tree. As a consequence, the length of the genealogical tree is proportional to the total number of polymorphic sites in the sample.

Genealogies under selection have long been an interesting object to study (see e.g. Wakeley, 2010 for a review). Starting with Krone and Neuhauser (1997) and Neuhauser and Krone (1997), genealogical trees under selection could be described using the Ancestral Selection Graph (ASG). In addition to coalescence events, which indicate joint ancestry of ancestral lines, selective events affect the genealogy in the following way: First, going backward in time, splitting events indicate possible ancestry. Since fit types are more likely to produce offspring in selective events, they are more likely to be true ancestors in such splitting events. However, true ancestry can only be decided once the ancestral types are known. So, in a second stage, going forward in time through the ASG, types are fixed and it can be decided which of the possible ancestors is true. The disadvantage of these splitting events is that they make this genealogical structure far more complicated to study than the coalescent for neutral evolution.

In recent years, much progress has been made in the simulation of genealogical trees under selection. Mostly, these simulation algorithms use the approach of the structured coalescent, which is based on Kaplan et al. (1988). Here, the allelic frequency path is generated first, and conditional on this path, coalescence events are carried out. (See also Barton et al., 2004 for a formal derivation of this approach.) Simulation approaches based on this idea include the inference method by Coop and Griffiths (2004), msms by Ewing and Hermisson (2010), and discoal Kern and Schrider (2016). However, the structured coalescent approach was only used in a few studies in order to obtain analytical insights (see e.g. Taylor, 2007).

Recently, genealogies under selection have been studied by Depperschmidt et al. (2012) using Markov processes taking values in the space of trees, i.e. the genealogical tree is modelled as a stochastic process which is changing as the population evolves. As for many Markov processes, the equilibrium can be studied using stationary solutions of differential equations. In our manuscript, we will make use of this approach in order to compute an approximation for the total tree length under a general bi-allelic selection scheme, which is assumed to be weak; see Section 4. Our results are extensions of Theorem 5 of Depperschmidt et al. (2012), where an approximation of the Laplace-transform of the genealogical distance of a pair of individuals under bi-allelic mutation and low levels of selection was computed.

The paper is structured as follows: In Section 2, we introduce the model we are going to study, i.e. genealogies in the large population limit for a Moran model under genic selection, incomplete dominance or over-or under-dominant selection. The last three cases we collect under the term *other modes of dominance*. We give recursions for the Laplace-transform (and the expectation) of the tree length of a sample in Theorem 1 and Corollary 4 for genic selection, and in Theorem 2 and Corollary 7 for other modes of dominance. In Section 3, we discuss our findings and also provide some plots, based on numerical solutions of the recursions, on the change of tree lengths under selection. Section 4 gives some preliminaries for the proofs. In particular, we give a brief review of the construction of evolving genealogies from Depperschmidt et al. (2012). Finally, Section 5 contains all proofs.

## 2 Model and main results

We will obtain approximations for the tree length under selection. While Theorem 1 and its corollaries describe the case of genic selection, Theorem 2 and its corollaries deal with other modes of dominance. All proofs are found in Section 5.

### Genic selection

Consider a Moran model of size *N*, where every individual has type either • or ∘, selection is genic, type • is advantageous with selection coefficient *α*, and mutation is bi-directional. In other words, consider a population of *N* (haploid) individuals with the following transitions:

1. Every pair of individuals resamples at rate 1; upon such a resampling event, one of the two individuals involved dies, the other one reproduces.
2. Every line is hit by a mutation event from ∘ to • at rate *θ*_∘_ > 0, and by a mutation event from • to ∘ at rate *θ*_•_ > 0.
3. Every line of type • places an offspring on a randomly chosen line at rate *α*.

Let *X* denote the frequency of • in the population. Mutation leads to an expected change *dX* of 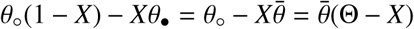 for 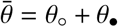 and 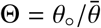 per time *dt*, and selection of *αX*(1 − *X*)*dt*. Recall that *X* follows – in the limit *N* → ∞ – the SDE

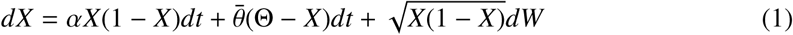

for some Brownian motion *W*; see e.g. (5.6) in Ewens (2004).

In the sequel, we will rely here on the possibility to study the genealogical tree of a sample taken in the large population limit of a Moran model in equilibrium. Since the time-point of sampling is the same for all ancestral lines, the resulting tree is ultra-metric and is given by genealogical distances between all pairs of individuals in the sample. In addition, marks on the tree describe mutation events from • to ∘ or from ∘ to •; see also Figure 1. This possibility is implicitly made by the ancestral selection graph from Neuhauser and Krone (1997), and formally justified by some results obtained in Depperschmidt et al. (2012); precisely, their Theorem 4 states that the genealogical tree under selection has a unique equilibrium.

**Figure 1:**
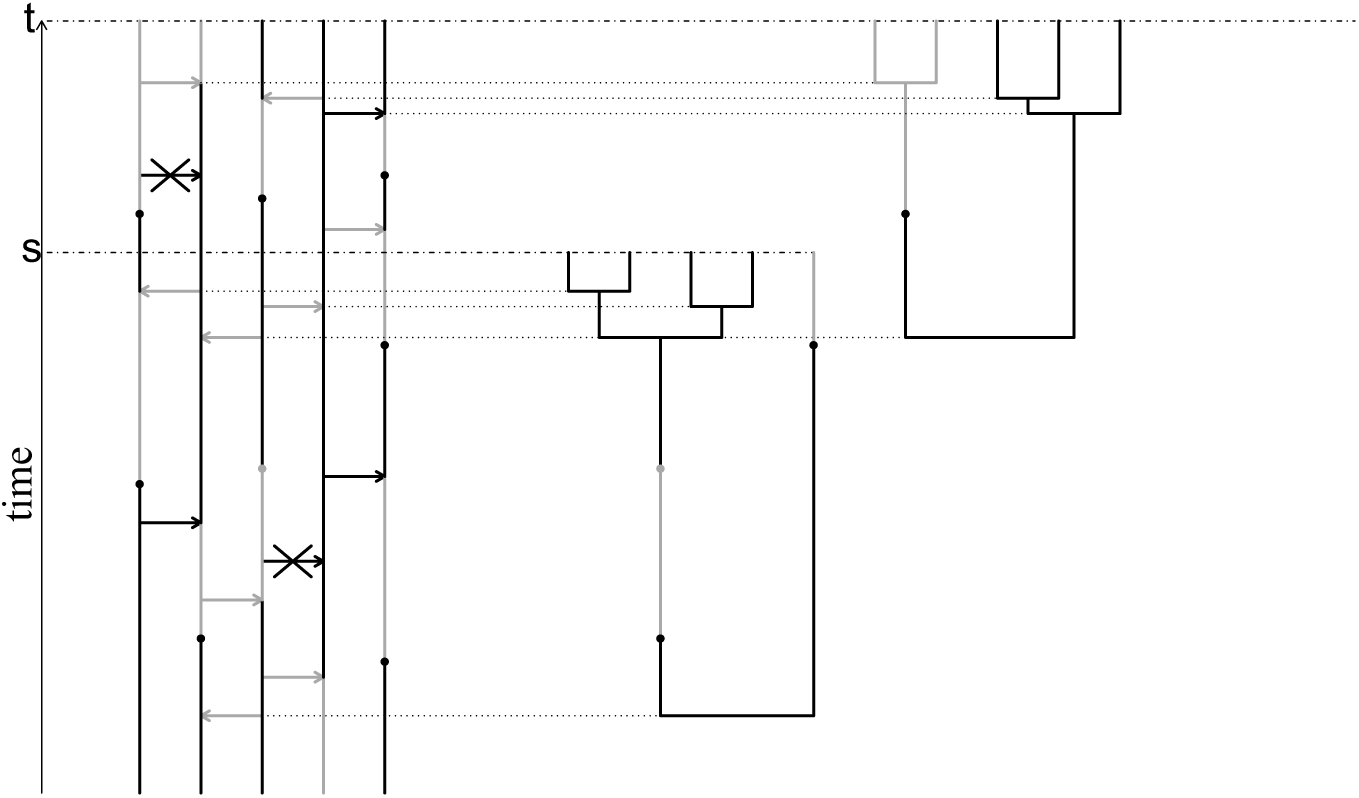
A population of size *N* = 5 following Moran dynamics with resampling, mutation, and genic selection in equilibrium. In the graphical representation on the left, bullet points indicate mutation events from the beneficial • type to ∘ and back. Grey arrows indicate neutral resampling events. Selective events are given through black arrows and only take effect if the arrows starts at the beneficial (black) type. Selective arrows starting with the deleterious type cannot be used and are indicated by a cross. Here, the full tree of all individuals in the model is drawn on the right for two points in time, but it is also possible to study the tree of a population sample.

We will write ℙ^*α*^[.] for the distribution of genealogical trees taken from the large population limit of Moran models under the selection coefficient *α* and 𝔼^*α*^[.] for the corresponding expectation. In particular, ℙ^0^[.] and 𝔼^0^[.] are reserved for neutral evolution, *α* = 0. Within the genealogical tree, we pick a sample of size *n* and let

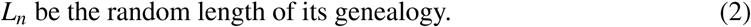

We note that in the absence of selection, *L*_*n*_ does not depend on the mutational mechanism and 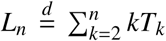, where 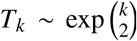, *k* = 2, …, *n* are the coalescence times in the tree; see e.g. (3.25) in Wakeley (2008). In particular, for *λ* ≥ 0,

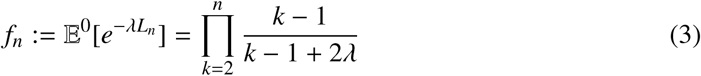

with *f*_1_ = 1 since the empty product is defined to be 1. We are now ready to state our first main result, which gives a recursion for an approximation of the Laplace-transform of the tree length under selection for small *α*.

#### Theorem 1

(Genealogical distances under genic selection). *Let* 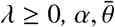 *and* Θ *be given as in* (1), *L*_*n*_ *as in* (2) *and*

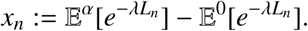

*Then, x*_1_, *x*_2_, … *satisfy the recursion x*_1_ = 0 *and*

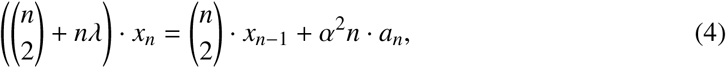

*where a*_1_, *a*_2_, … *satisfy a recursion a*_1_ = 0 *and*

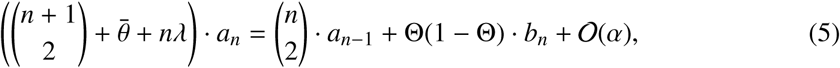

*where b*_1_, *b*_2_, … *satisfy a recursion b*_1_ = 0 *and*

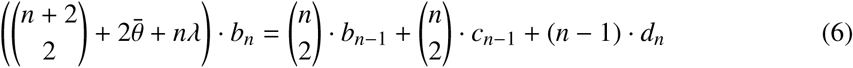

*where c*_1_, *c*_2_, … *satisfy a recursion c*_1_ = 0 *and*

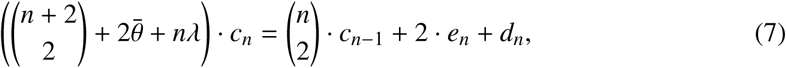

*where e*_1_, *e*_2_, … *satisfy a recursion e*_1_ = 0 *and*

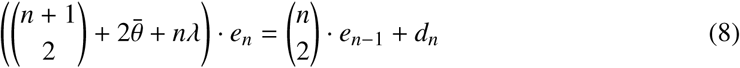

*and finally – recall* (3) *–*

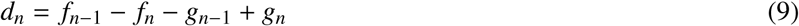

*with* 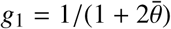 *and*

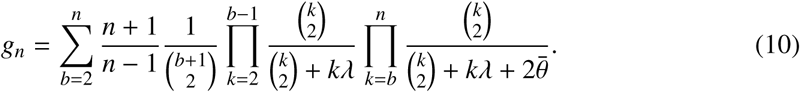

#### Remark 1

(Interpretations). In the proof, we will see that the quantities *a*_*n*_, *b*_*n*_, *c*_*n*_, … do have interpretations within the Moran model. If the tree length of the genealogy of a sample of individuals 1, …, *n* is denoted *L*_*n*_, the genealogical distance of individuals *i* and *j* is *R*_*ij*_, and *U*_*i*_ is the type of individual *i* (either • of ∘), these are

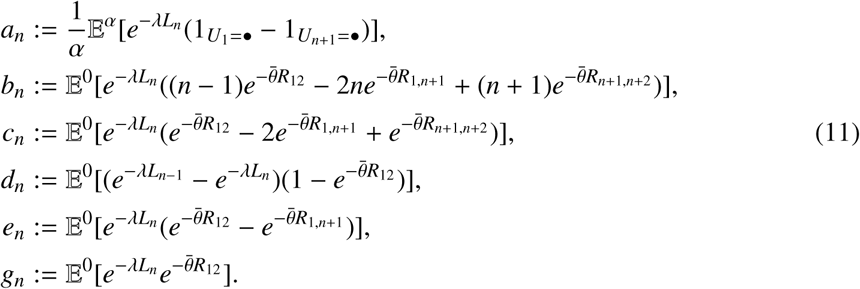

Moreover, in Theorem 2, another quantity will arise, which is

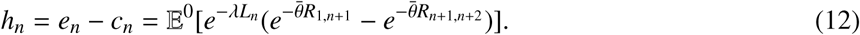

We note that, from these definitions, only *a*_*n*_ relies on the model with selection (since all other terms are within the neutral model. In addition, *a*_1_ = *b*_1_ = *c*_1_ = *e*_1_ = 0. The initial value *d*_2_ is given through the initial condition *f*_1_, as well as *f*_2_, *g*_1_ and *g*_2_.

#### Remark 2

(Comparing neutral and selective genealogies).
1. Note that for *α* = 0, (4) gives precisely (3). Moreover, there is no linear term in *α* in the recursion (4). This finding is reminiscent of Theorem 4.26 in Krone and Neuhauser (1997) and Theorem 5 in Depperschmidt et al. (2012), but we note that for other models of dominance, a linear term arises; see Theorem 2.
2. Let us compare tree lengths under neutrality and under selection qualitatively. Crucially, the quantity *d*_*n*_ as given in (11) is positive. As consequences, by the recursions, *e*_*n*_ from (8) is positive, *c*_*n*_ from (7) is positive, *b*_*n*_ from (6) is positive, and *a*_*n*_ from (5) is positive. The effect is that *x*_*n*_ for small *α* is positive, i.e. 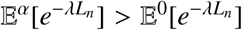 for small *α*, which implies that genealogical trees are generally shorter (in the so-called Laplace-transform-order) under selection. In particular, we have shown the intuitive result that expected tree lengths are shorter under selection; see also Corollary 4 for a quantitative result concerning expected tree lengths.
3. While *x*_*n*_, *a*_*n*_ are quantities within the selected genealogies, all other quantities can be computed under neutrality, *α* = 0. However, if one would like to obtain finer results, i.e. specify the 𝒪 (*α*^3^)-term in (4), more quantities within selected genealogies would have to be computed. In principle, this is straight-forward using our approach of the proof of Theorem 1.

#### Remark 3

(Solving the recursions). All recursions for *x*_*n*_, *a*_*n*_, *b*_*n*_, *c*_*n*_, *e*_*n*_, *h*_*n*_ are of the form

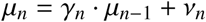

with *µ*_1_ = 0 and can readily be solved by writing

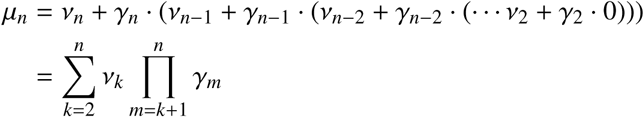

with Π_∅_ := 1.

Since we can directly obtain expected tree lengths from the Laplace-transforms in Theorem 1, we obtain also a recursion for expected tree lengths by using that 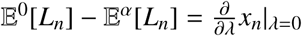.

#### Corollary 4

(Expected tree length under genic selection). *With* 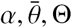 *and L*_*n*_ *as in Theorem 1, let*

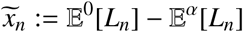

*Then,* 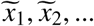 *satisfy the recursion* 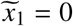 *and*

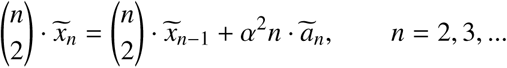

*where* 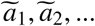 *satisfy a recursion* 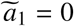 *and*

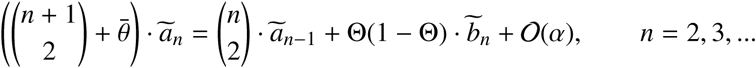

*where,* 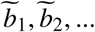 *satisfy the recursion* 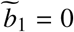 *and*

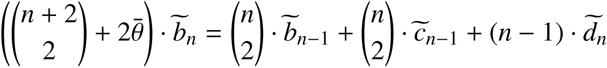

*where,* 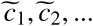 *satisfy the recursion* 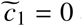 *and*

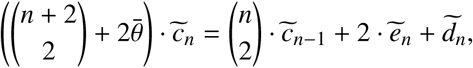

*where,* 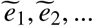 *satisfy the recursion* 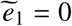 *and*

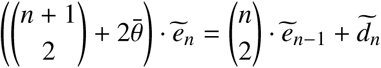

*and finally*

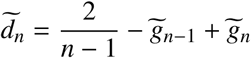

*with* 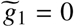 *and*

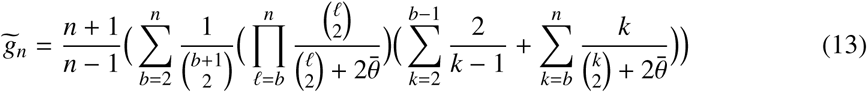

The following result, the special case *n* = 2, was already obtained in Theorem 5 of Depperschmidt et al. (2012).

#### Corollary 5

(Genealogical distance of two individuals under genic selection). *With* 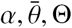 *and L*_*n*_ *as in Theorem 2,*

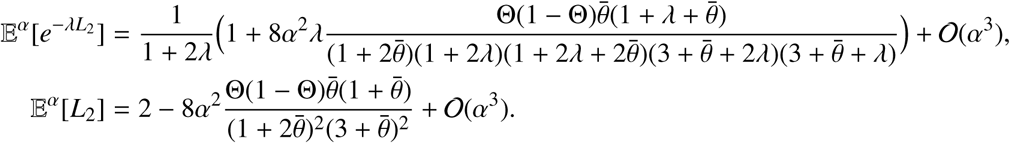

### Other modes of dominance

In a diploid population, (1) only models the frequency of • correctly if selection is genic, i.e. if the selective advantage of an individual which is homozygous for • is twice the advantage of a heterozygote. For other modes of dominance, we have to introduce a dominance coefficient *h*∈(–∞,∞) and change the dynamics of the Moran model as follows: Let *X* be the frequency of •in the population. In addition to 1. and 2. from the beginning of Section 2, we add frequency-dependent selection events:

3’. Every line of type • places an offspring to a randomly chosen line at rate *α*(*X* + *h*(1 − *X*)). Every line of type ∘ places an offspring to a randomly chosen line at rate *αhX*.

Note that 3’. is best understood by assuming that every line picks a random partner and if the pair is a heterozygote, it has fitness advantage *αh*, and if it is homozygous for •, it has fitness advantage *α*. (Here, we have assumed that *h* ≥ 0, but some modifications of 3’. also allow for *h* < 0.) The expected effect of 3’. on *X* is then *αX*(1 − *X*)(*X* + *h*(1 − *X*)) − *αX*(1 − *X*)*hX* = *αX*(1 − *X*)(*h*+(1 − 2*h*)*X*) and the frequency of • follows – in the limit *N* → ∞ – the SDE

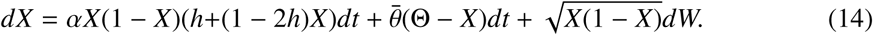

We will write ℙ^*α,h*^[.] for the distribution of genealogical trees and allele frequencies under this scenario, and 𝔼^*α,h*^[.] for the corresponding expectation. Recalling that ℙ^*α*^[.] and 𝔼^*α*^[.] are the correpsonding operators for genic selection, we have ℙ^*α*^[.] = ℙ^2*α*,1/2^[.]. We note that 3’. above for 2*α* and *h* = 1/2 does not directly transform to 3. under genic selection for selection intensity *α*. However, as argued above (14), the effect on the process *X* only comes from the difference of the effects of type • and ∘ and the same argument applies on the level of genealogies, and this difference is the same for 3. and 3’.

We note that *h* = 0 means a positively selected recessive allele, while *h* = 1 refers to a dominant selectively favoured allele. Again, we obtain an approximation of the Laplace-transform of the tree length of a sample of size *n*.

#### Theorem 2

(Genealogical distances under any form of dominance). *Let* 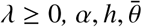 *and* Θ *be given as in* (14), *L*_*n*_ *as in* (2) *and*

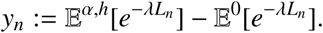

*Then, y*_1_, *y*_2_, … *satisfy the recursion y*_1_ = 0 *and*

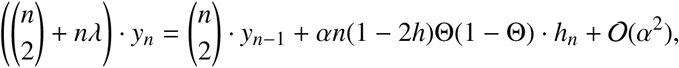

*where h*_1_, *h*_2_, … *satisfy the recursion h*_1_ = 0 *and*

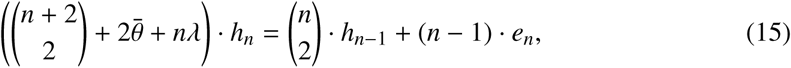

*and e*_*n*_ *was given in Theorem 1.*

#### Remark 6

(Comparing genealogies).
1. Most interestingly, neutral trees differ from trees under genic selection only in order *α*^2^, whereas the difference is in order *α* for other forms of dominance. While this may be counter-intuitive at first sight, it can be easily explained. Note that the model actually does not change if we replace *α* by −*α* and *h* by 1 − *h* at the same time. By doing so, we just interchange the roles of allele • and ∘. For *h* = 1/2, this means that our results have to be identical for *α* and −*α*, leading to a vanishing linear term in (4). For *h* ≠ 1/2, this symmetry does not have to hold, leading to a linear term in *α*.
2. Similar to our reasoning in Remark 2.2, the sign of *h*_*n*_ in the recursion for *y*_*n*_ determines if tree lengths are shorter or longer under selection. We see that the behaviour changes at *h* = 1/2. By construction, *h*_*n*_ is positive, so if *h* < 1/2, *y*_*n*_ is positive as well and we see that trees are shorter under selection (in the Laplace-transform order). If *h* > 1/2, the reverse is true and trees are longer under selection. This result is not surprising for over-dominant selection, *h* > 1, since the advantage of the heterozygote leads to maintenance of heterozygosity or balancing selection, which in turn is known to produce longer genealogical trees.

#### Corollary 7

(Expected tree length under any form of dominance). *With* 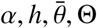 *and L*_*n*_ *as in Theorem 2, let*

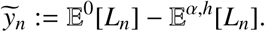

*Then,*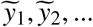 *satisfy the recursion* 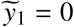 *and*

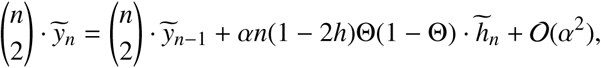

*where* 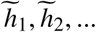 *satisfy the recursion* 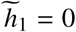 *and*

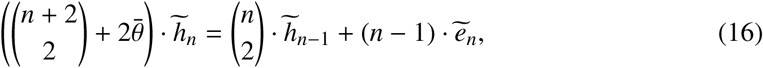

*and* 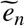 *was given in Corollary 4.*

#### Corollary 8

(Genealogical distance of two individuals under any form of dominance). *With* 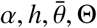 *and L*_*n*_ *as in Theorem 2,*

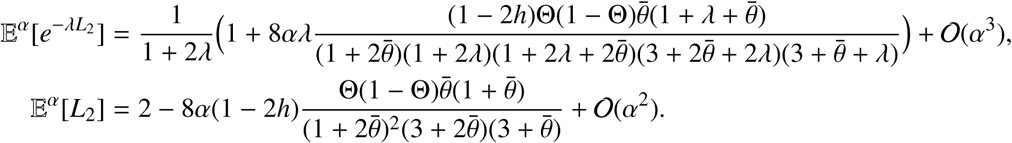

## 3 Discussion

A fundamental question in population genetics is: How does selection affect genealogies of a sample of individuals? We have added to this question an analysis of tree lengths under low levels of selection, both for genic selection and for other modes of dominance. While our results are only given through recursions, these give valuable insights. Recall that under neutrality,

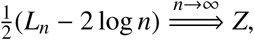

where *Z* is Gumbel distributed (see p. 255 of Wiuf and Hein, 1999). In particular, for large *n*,

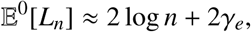

where *γ*_*e*_ ≈ 0.57 is the Euler-Mascheroni constant.

Summing up and extending previous results for large samples, the main findings of our study are the following on the expected tree lengths under selection. A proof is found in the appendix.

### Proposition 9

(Summary). *For h* = 1/2, *there is C*_1/2_ < ∞ *such that*

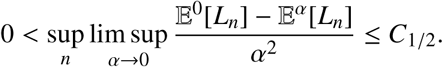

*For h* < 1/2, *there is C*_<1/2_ < ∞ *such that*

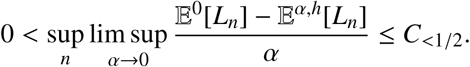

*For h* > 1/2, *there is C*_>1/2_ < ∞ *such that*

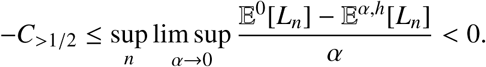

Since 𝔼^0^[*L*_*n*_] = 𝒪 (log *n*) for large *n*, this shows that the order of magnitude in the change due to selection is much smaller than the length of the full tree for large samples. Note that this finding is in line with Przeworski et al. (1999), where simulations of the ancestral selection graph are used to find that the overall tree shape under selection is not too different from neutrality for low levels of selection.

In order to get more quantitative insights, we have numerically solved the recursions from Corollaries 4 and 7, and plotted the effect on the tree length for various scenarios. Figure 2(A) analyses the effect of genic selection in large samples (i.e. *n* = 50). Interestingly, there is some mutation rate 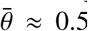, which gives the largest effect. This is clear since very little mutation implies almost no change in the genealogy relative to neutrality since almost always only the beneficial type is present in the population, and very high mutation rate implies that selection is virtually inefficient, leading to nearly neutral trees. Moreover, we can see here that Θ(1 − Θ) enters the recursion for the change in tree length only linearly. In Figure 2(B), we display the change in tree length for *h* = 0. Since 1 − 2*h* enters the recursion only linearly, the graph looks qualitatively the same for other dominance coefficients.

**Figure 2:**
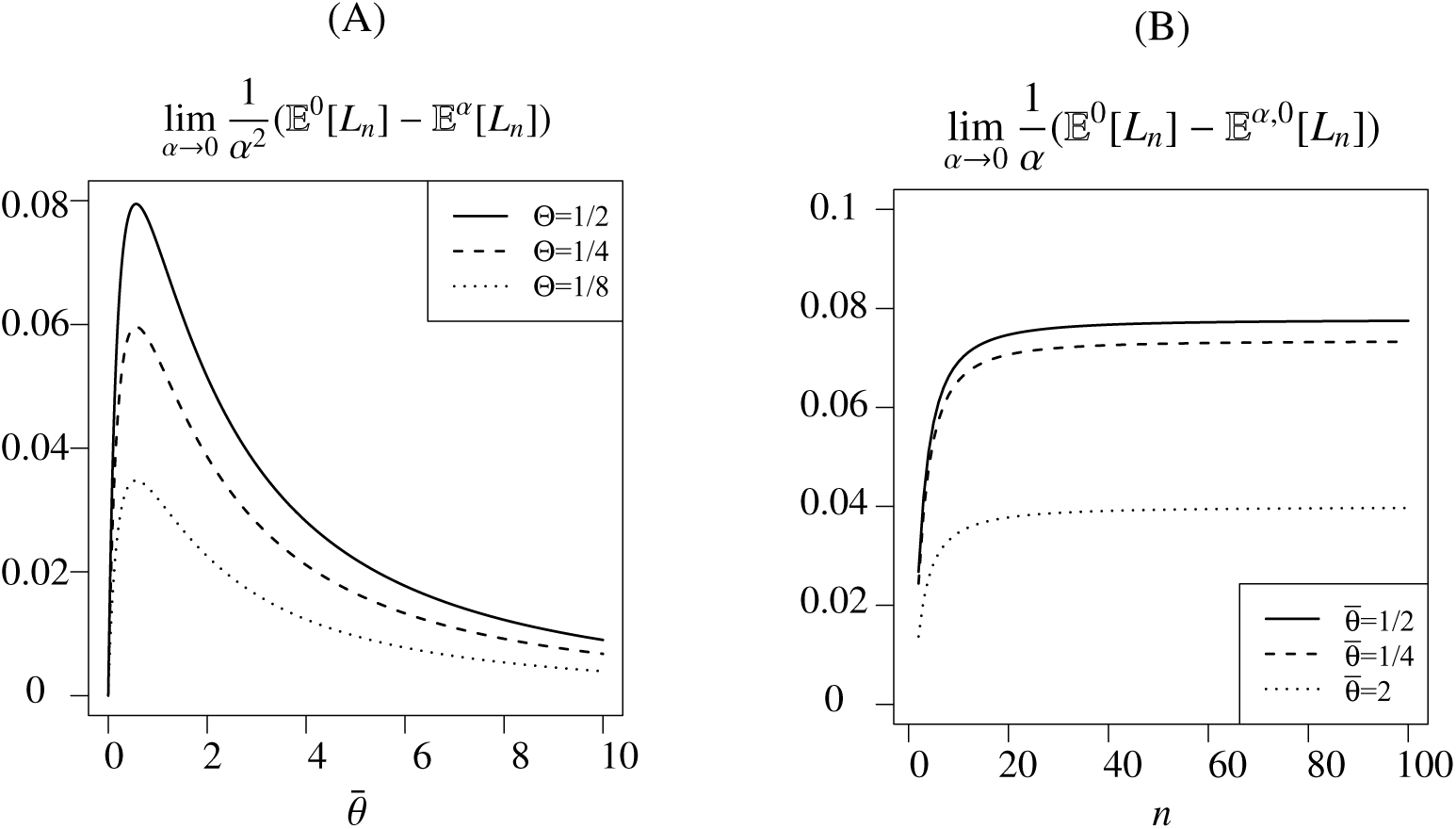
Using the recursions from Corollaries 4 and 7, we see differences in expected tree length. (A) For genic selection and large samples, the effect changes with the total mutation rate 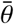 and is linear in Θ(1 – Θ). (B) Plot of the change in total tree length for small values of *α* with *h* = 0, dependent on the sample size, and three parameters of 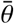.

In principle, the approach we use here is comparable to the ancestral selection graph in the sense that all events happening within the ASG are also implemented in our construction. However, within the ASG, when a splitting event occurs, it is not clear which of the two lines is the true ancestor, so both lines are followed. As a consequence, in any case the ASG *looks* longer than a neutral coalescent due to the splitting events, and only when the ASG is pruned to become the true genealogical tree, can the tree length be computed. In our approach, the information, which we need within a splitting event is different, since only the type of the additional line, and the type of an individual within the sample is needed.

The approach we use here to study genealogies under selection can be used for other statistics than the total tree length. In principle, every quantity of a sample tree can be described. The reason why we chose the tree length is its simple structure due to coalescence events: If two lines in a sample of size *n* coalesce, all that remains is a tree of *n* – 1 individuals, already implying the recursive structure for the tree lengths which is apparent in Theorems 1 and 2. In addition, in order to describe the effect of selection, we rely on a description of the sample which was already used in the mathematics literature for the so-called Fleming-Viot process (which generalizes the Wright-Fisher diffusion); see e.g. Etheridge (2001) and Depperschmidt et al. (2012). In principle, our approach can be extended e.g. to include more than two alleles, population structure, recombination etc. However, one would still have to find a recursive structure, which will often be feasible only in the case of weak selection.

## 4 Preliminaries for the proofs

We here present the construction of a tree-valued process in a nutshell, leaving out various technical details. All details of the construction are given in Depperschmidt et al. (2012).

Any genealogical tree is uniquely given by all genealogical distances between pairs of (haploid) individuals. So, in order to describe the evolution of genealogical trees, it suffices to describe the evolution of all pairwise distances. We also note that the tree length which we consider in all our results, is a function of pairwise distances. (See e.g. Section 8 of Depperschmidt et al. (2012).) Consider a sample of size *n* taken from the Moran model at time *t* as described in Section 2, and let *R*(*t*) := (*R*_*ij*_(*t*))_*i*<*j*_ be the pairwise genealogical distances (note that *R*_*ij*_ = *R* _*ji*_ and *R*_*ii*_ = 0), and *U*(*t*) := (*U*_1_(*t*), …, *U*_*n*_(*t*)) the allelic types, either •or ∘. Within the sample of size *n*, we will speak of *R*(*t*) as the sample tree, and of *U*(*t*) of the types within the sample. Note that there is the possibility to extend the sample by picking new individuals at random, leading to types *U*_*n*+1_(*t*), *U*_*n*+2_(*t*), …, which we will need for selective events below. We will consider some smooth, bounded function: 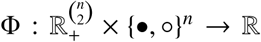 and are going to describe the change in 𝔼^*α,h*^[Φ(*R*(*t*), *U*(*t*))] due to the evolution of the Moran model. We have to take into account several mechanisms:

0. Growth of the tree: During times when no events happen, all genealogical distances grow deterministically and linearly (with speed 2). In time *dt*, the change is, using the partial derivative of Φ with respect to the *i j*th coordinate in 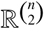, denoted by 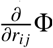,

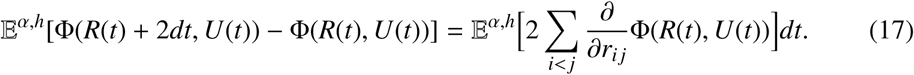
1. Resampling: If a pair of the *N* individuals within the Moran model resamples, there is either none, one, or two of them within the sample of size *n*. If none, the sample tree is not affected. If one, and this one reproduces, the sample tree is not affected as well. If one, and this one is replaced by the individual outside of the sample, the effect is the same as if we would have picked the other individual to begin with. Since in the 𝔼^*α,h*^[…], we average over all possibilities which samples of size *n* we take, there is also no resulting effect. If two, *i* reproduces and *j* dies, say, the effect is that distances to individual *j* are replaced by distances to individual *i*, and the new type of individual *j* is the type of *i*. Since all pairs within the sample resample at rate 1, the change in time *dt* is

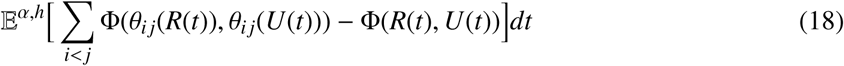

With

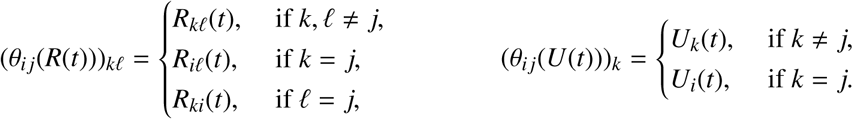
2. Mutation: We note that the mutational mechanism can also be described by saying that every line in the Moran model mutates at rate 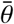, with outcome • with probability Θ and outcome ∘ with probability 1– Θ. Since mutation only affects the alleles of the individuals in the sample, we have in time *dt*

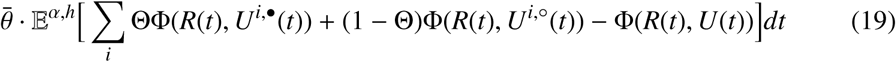

With

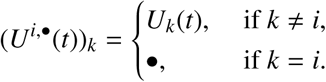

and analogously for *U*^*i*,∘^.
3. Selection: By the dynamics of the Moran model, we have to accept that selective events depend on the type of individuals. We say that the *k*th individual has fitness *α*χ_*k*_ := *α*χ(*U*_*k*_), and χ is the fitness function. For genic selection and other modes of dominance, the fitness functions are denoted by χ_*k*_ and χ_*k,m*_, respectively, where for other modes of dominance, *m* is some randomly picked (haploid) individual. (Note that *n* « *N*, so we have that *m* is outside of the sample with high probability.) The fitness functions are given by

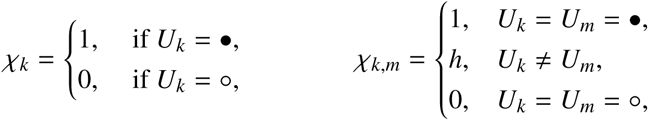

respectively, where we will have 1 ≤ *k* ≤ *n* and *m* > *n* below. As for resampling, selective events occur in a pair of one individual *i* giving birth and the other, individual *j*, dying, as given through the function *θ*_*ij*_ from above. We start with genic selection (recall that 𝔼^*α*^[.] = 𝔼^2*α*,1/2^[.]). Here, we find that in time *dt*, since *n* « *N*,

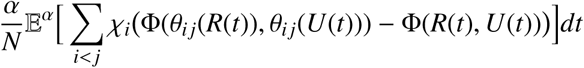

where summands are zero, if *j* > *n*, and summands with *i ≤ n* only give a negligible effect; for the remaining summands, *i* is any individual outside of the sample, so we choose *i* = *n* + 1 without loss of generality and obtain

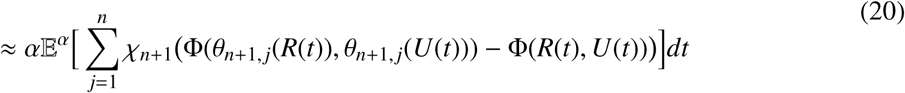

which gives by permuting sampling order of *j* and *n* + 1 in the first term,

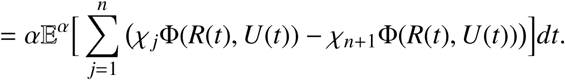

Note that the ≈ is exact in the limit *N* → ∞. For other modes of dominance, we find analogously the effect

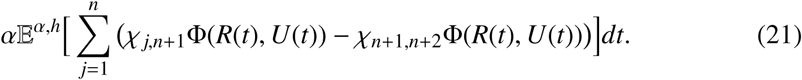

In the calculations above, Φ can be any smooth function, which depends on a randomly picked sample. Now, we consider a sample of size *n* + *j* for some *n, j* ≥ 0 and focus on the function, for some 0 ≤ *i* ≤ *n*,

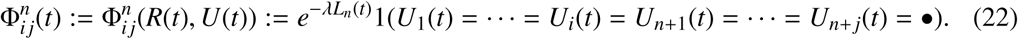

Note that although 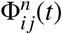 is dealing with a sample of size *n* + *j*, the tree length is only computed with respect to the genealogical tree of the first *n* individuals. In words, the quantity 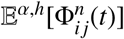 is the Laplace-transform of the length of the genealogical tree of a sample of size *n* under selection, on the event that the first *i* individuals within the sample, as well as *j* additionally picked individuals (outside of the sample) carry the beneficial type •. For *i* = *j* = 0,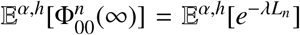 is thus the Laplace-transform of the tree length of a sample of size *n* under selection and the main object of study in Theorems 1 and 2. The following lemma is an application of the general theory from 0.-3. above.

### Lemma 10

*For n* ≥ 2, *and with the convention that* 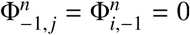, *in the limit N* → ∞,

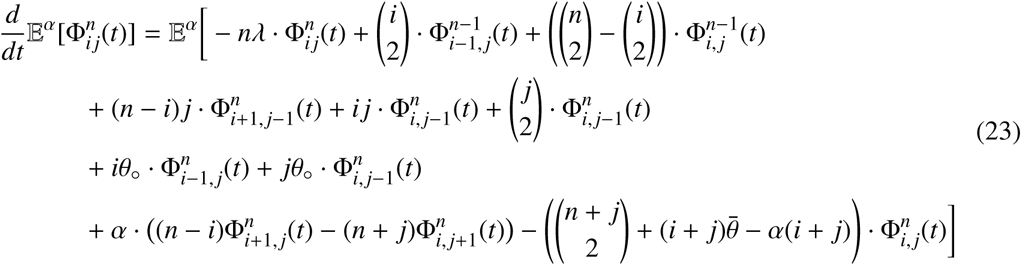

*for genic selection, and the last two terms change to*

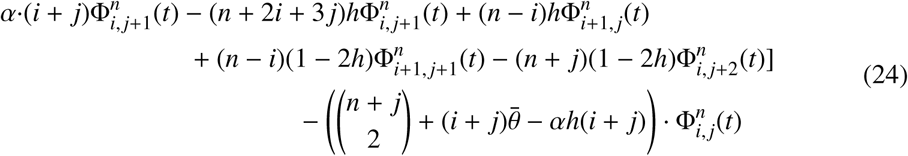

*for other modes of dominance.*

### Remark 11

Simple algebra shows that the *α*-terms in (23) and (24) agree for *h* = 1/2, i.e. (23) with 𝔼^*α*^[.] agrees with (24) for 𝔼^2*α*,1/2^[.] For future reference, note that for *i* = *j* = 0, the *α*-term in (24) gives

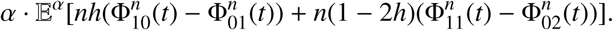

*Proof of Lemma 10.* We will omit dependencies on *t* and write 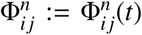 in the proof. The effect of tree growth on 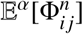 is that the tree grows by *ndt* in time *dt*, i.e. in time *dt*

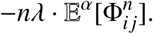

Let *I* = {1, …, *i*}, *H* = {*i* + 1, …, *n*} and *J* = {*n* + 1, …, *n* + *j*}. For resampling, we distinguish between events among *I*, events among *I* ∪ *H*, with at most one partner within *I*, events with one partner within *I* and the second among *J*, events with one partner in *H* and the second among *J*, and events with two partners in *J*. Only if two among *I* ∪ *H* coalesce, *n* decreases. This gives

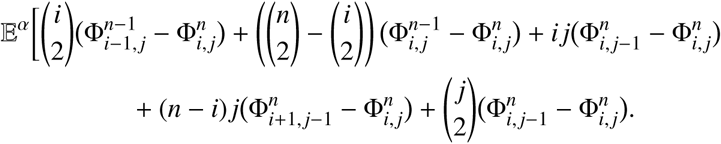

For mutation, we note that for 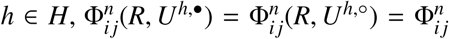, herefore the effect of mutation is

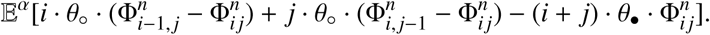

Last, for selection, we have to distinguish the cases of genic selection and other modes of dominance. For genic selection, we have that χ_*k*_ = 1(*U*_*k*_ = •) and we note that for *i* ∈ *I* and *j* ∈ *J*,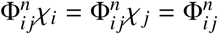, therefore the effect is, from (20),

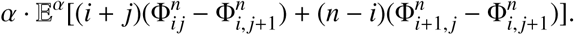

For other modes of dominance

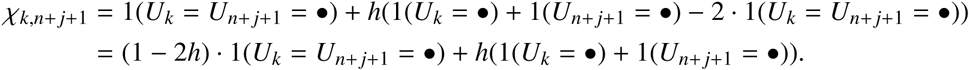

Therefore, the effect of selection is here, from (21),

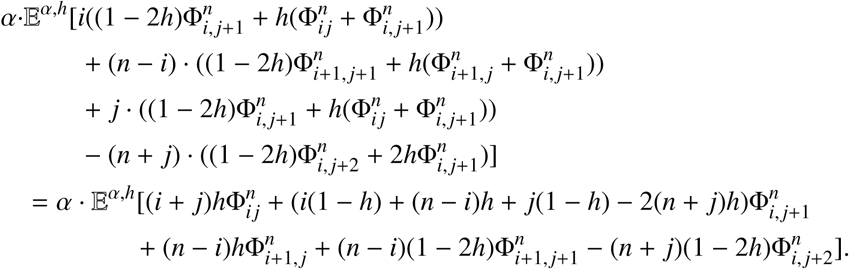

□

The next result is collected from Theorem 4 and Lemma 8.1 in Depperschmidt et al. (2012).

### Lemma 12

*The process* (*R, U*) *of genealogical distances and types, has a unique equilibrium under* ℙ^*α,h*^. *This equilibrium is described by* 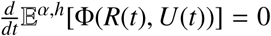 *for all possible* Φ. *Moreover, for this equilibrium, denoted by* (*R*(∞), *U*(∞)), *satisfies*

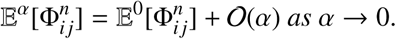

## 5 Proof of Theorems 1 and 2

*Proof of Theorem 1.* To begin, we note that the quantities as defined in Remark 1 for *n* = 1 – since *L*_1_ = 0 – are given by *a*_1_ = *b*_1_ = *c*_1_ = *e*_1_ = 0. Moreover, we note that 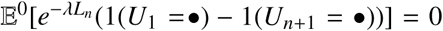 since the mutational history of single lines, leading to *U*_1_ and *U*_*n*+1_ are independent of the genealogy for *α* = 0 and therefore, from Lemma 12, we see that *ã*_*n*_ := *αa*_*n*_ = 𝒪 (*α*). For the recursion on *x*_*n*_, we write – from Lemma 10 –

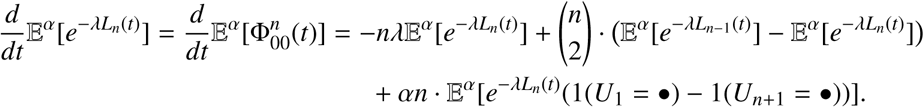

In equilibrium, the right hand side must equal 0, and using this equality also for *α* = 0, we have

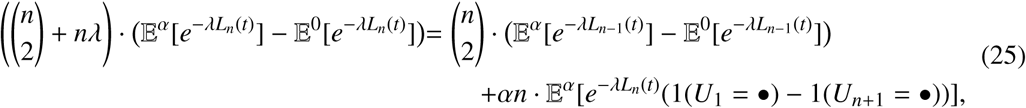

which gives exactly (4), where *a*_*n*_ is given in (11). For the recursion on *a*_*n*_, we write with Lemma 10

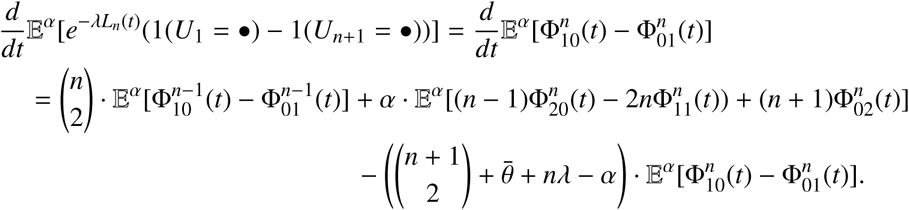

In equilibrium, both sides must be 0, and dividing both sides by *α* gives a recursion for *a*_*n*_, but we still have to show that the last term is *b*_*n*_ + 𝒪 (*α*). First, we can replace 𝔼^*α*^[.] by 𝔼^0^[.] in this expression, since we are making an error of order *α* at most. Then, for *α* = 0, i.e. neutral evolution, we note that two individuals *k* ≠ 𝓁, which have genealogical distance *R*_*k*𝓁_, both have type • in either of two cases: (i) there is no mutation on the path between *k*, 𝓁 in the genealogy, and their joint ancestor has type •; (ii) there is a mutation on the path between *k*,𝓁, and both mutational events determining the types of *k*, 𝓁 give the type •. For *α* = 0, the mutational process is independent of coalescence events, hence we find the probabilities 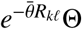 and 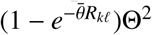 in cases (i) and (ii), respectively. Hence, for any *k*, 𝓁 = 1, …, *n* + 2 and *k* ≠ 𝓁

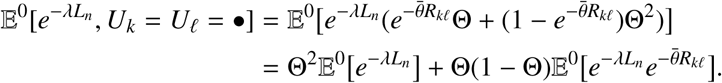

Hence, we obtain that

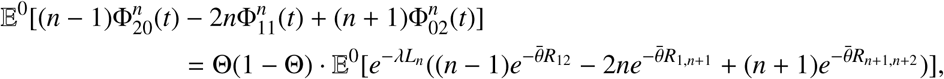

which proves (5), where *b*_*n*_ is given as in (11). For the remaining recursions, we always work with *α* = 0 and set 𝔼[.] := 𝔼^0^[.]. In order to obtain a recursion for *b*_*n*_, consider a coalescent with *n* + 2 lines and distinguish the following cases for the first step:

1. Coalescence of lines among the first *n* lines, except for lines 1,2 (rate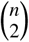 − 1);
2. Coalescence of lines 1,2 (rate 1);
3. Coalescence of lines *n* + 1 and 1 (rate 1);
4. Coalescence of lines *n* + 1 and one of 2, …, *n* (rate *n* − 1);
5. Coalescence of lines *n* + 1 and *n* + 2 (rate 1);
6. Coalescence of lines *n* + 2 and one of 1, …, *n* (rate *n*);

Recalling 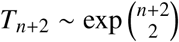, we write by a first-step decomposition

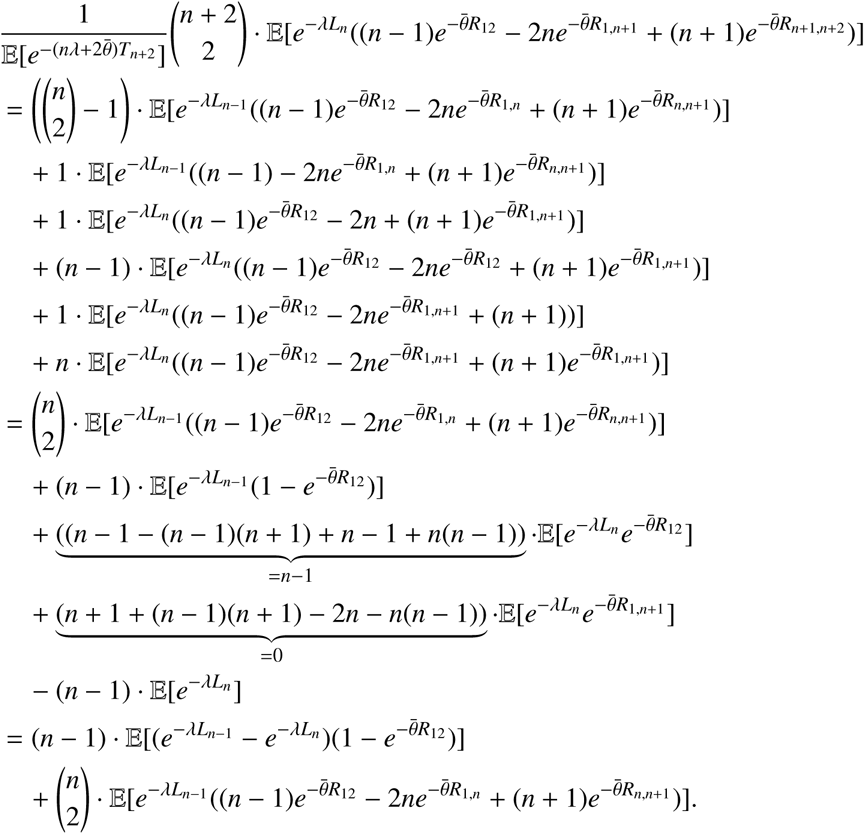

This shows (6). For *c*_*n*_, we use the same coalescent, and by distinguishing the six cases, we write

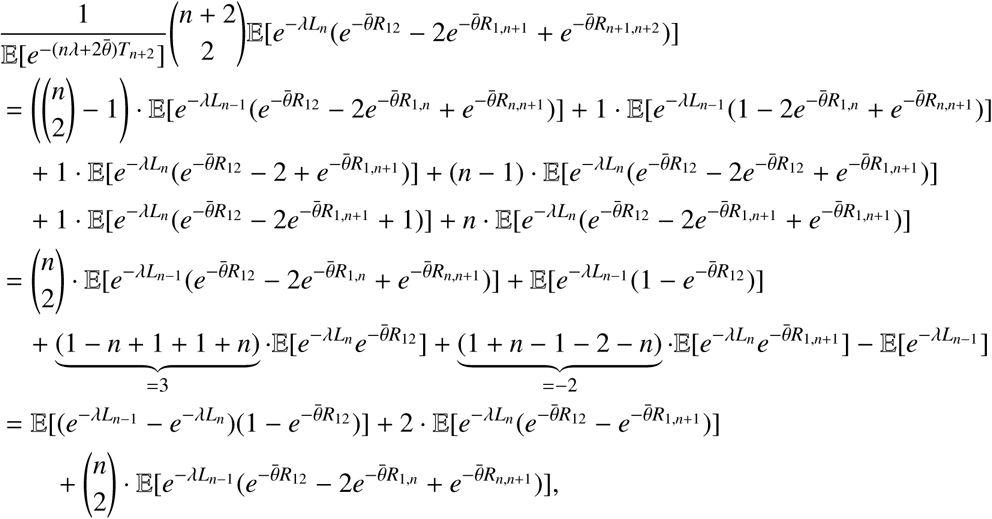

which shows (7). For *d*_*n*_, let *B* ∈ {2, …, *n*} be the number of lines in a coalescent starting with *n* lines, just before lines 1 and 2 coalesce. Then,

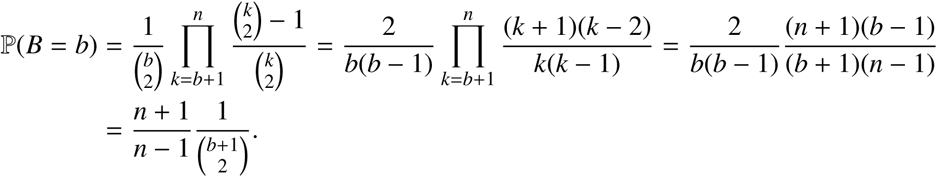

Then,

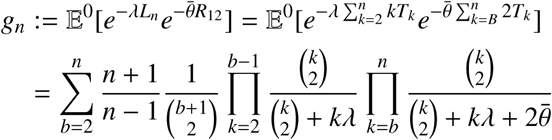

and we see that

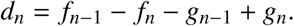

Finally, for *e*_*n*_, we again use a recursion. Consider a coalescent with *n* + 1 lines and make a first-step-analysis. In this first step, we distinguish four cases:

1. Coalescence of lines 1 or 2 with one of 3, …, *n*; rate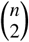 − 1
2. Coalescence of lines 1 and 2; rate 1
3. Coalescence of lines *n* + 1 and 1; rate 1
4. Coalescence of lines *n* + 1 and one of 2, …, *n*; rate *n* − 1

Hence,

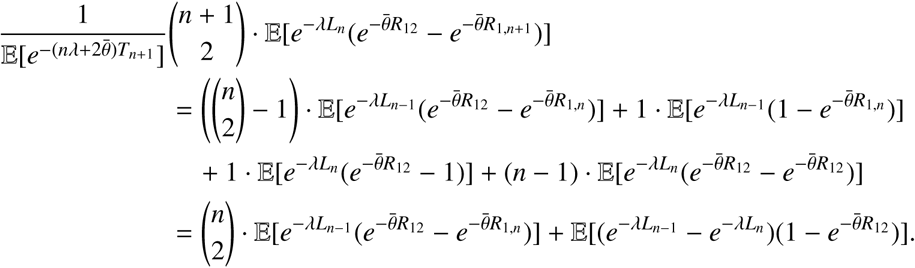

This shows (8).□

*Proof of Corollary 4.* We have to compute 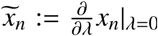. A close inspection of the recursions for *x*_n_ reveals that (i) *x*_*n*_ is a sum of products, and in each summand, some factor *d*_*k*_ enters and (ii) *d*_*k*_ = 𝒪 (λ) for small λ for all *k*, which is best seen from (11). As a consequence, we can compute the derivative with respect to λ at λ = 0 in each summand which enters *x*_*n*_ by taking the derivative of *d*_*k*_ with respect to λ and set λ = 0 in all other factors. Summing up, we have the same recursions as in Theorem 1 with (i) λ = 0 in all terms except *d*_*n*_, and (ii) replace *d*_*n*_ with the derivative according to λ at λ = 0. This gives the recursions as given in the corollary and

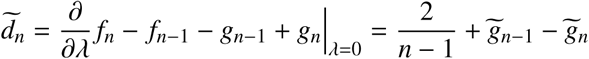

with

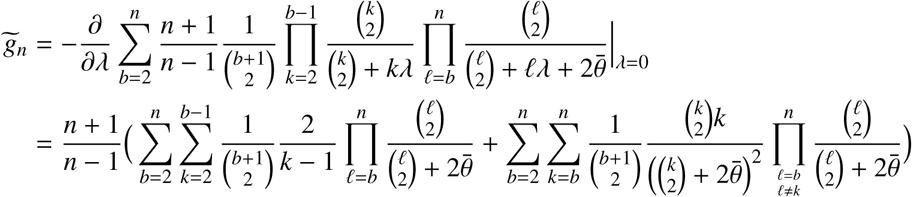

and the result follows. □

*Proof of Corollary 5.* Applying Theorem 1, we get

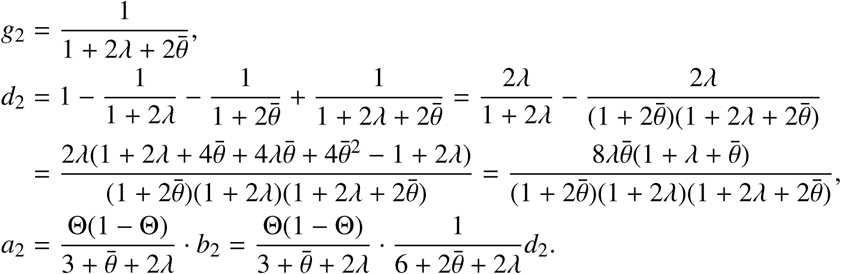

This gives the first assertion. The second follows since *a*_2_ is 𝒪 (*λ*) and the derivative with respect to *λ* at *λ* = 0 is easily computed. □

*Proof of Theorem 2.* Starting in the same way as in the proof of Theorem 1, we get, as in (25) – recall (20) –

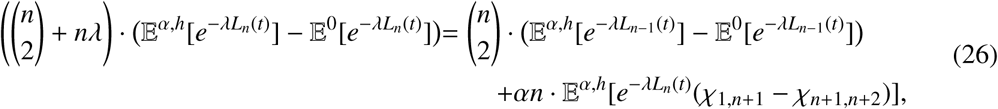

For the last term, we compute

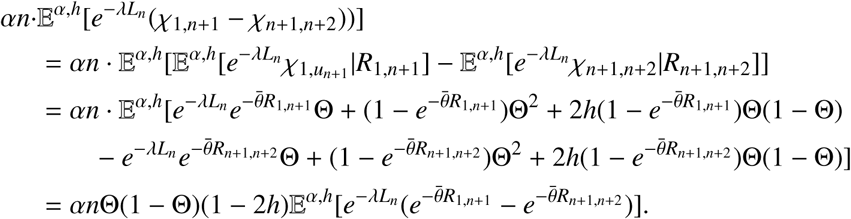

Now, we use that 𝔼^*α,h*^[.] = 𝔼^0^[.] + 𝒪 (*α*) as in Lemma 12 in order to obtain an approximate recursion for the last term. Consider a coalescent with *n* + 2 lines and distinguish the following cases:

1. Coalescence of lines among the first *n* lines (rate 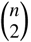);
2. Coalescence of lines *n* + 1 and 1 (rate 1);
3. Coalescence of lines *n* + 1 and one of 2, …, *n* (rate *n* − 1);
4. Coalescence of lines *n* + 1 and *n* + 2 (rate 1);
5. Coalescence of lines *n* + 2 and one of 1, …, *n* (rate *n*).

We then get

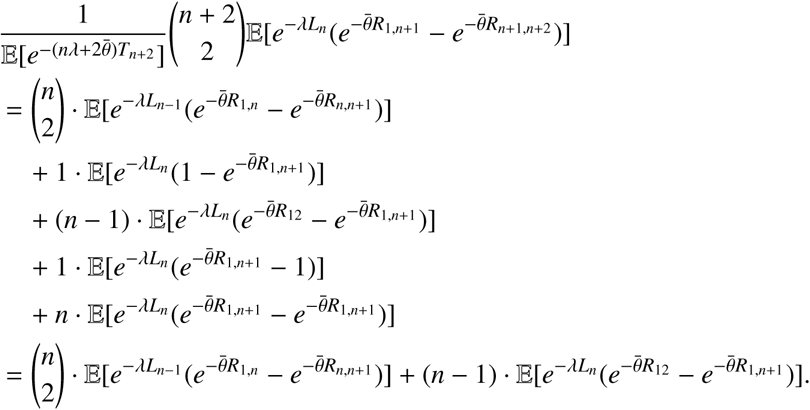

Using the definitions of *h*_*n*_ and *e*_*n*_ from Remark 1 then gives the result. □

*Proof of Corollary 7.* The proof is basically the same as for Corollary 4: The recursions for *y*_*n*_ is a sum of products, where each factor comes with a factor *d*_*n*_, and *d*_*n*_ = 𝒪 (*λ*). Therefore, the derivative according to *λ* at *λ* = 0 is performed by taking derivatives only of *d*_*n*_ and setting *λ* = 0 in all other instances. □

*Proof of Corollary 8.* Applying Theorem 2, we get

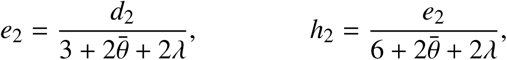

and with *d*_2_ from the proof of Corollary 5, the result follows. □

## A Proof of Proposition 9

For finite *n*, all results can be read off from Corollaries 4 and 5; see also Remark 6.1 for genic selection and Remark 6.2 for other modes of dominance. It remains to show uniformity in *n*. For the first assertion, we have that 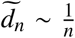, where *a*_*n*_ ∼ *b*_*n*_ if 0 < lim inf 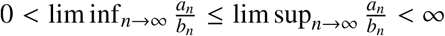. From Remark 3, we can solve the recursions for 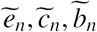 and *ã*_*n*_, which all are of the form

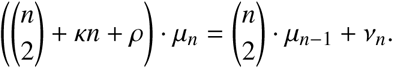

We want to find the behaviour of *µ*_*n*_ for large *n*. Hence,

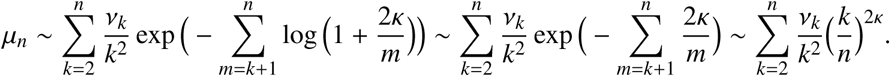

Since 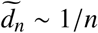 and the recursion for 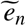 comes with *κ* = 1, we find that

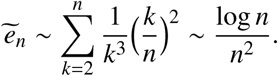

Next, the recursion for 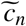 comes with *κ* = 2, and 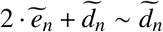, so 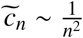. In the recursion for 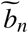, we have *κ* = 2, so 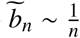. In the recursion for *ã*_*n*_, we have *κ* = 1, so 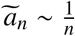 log *n*. Finally, for 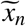, we write

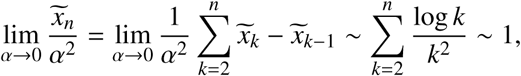

the first assertion of Proposition 9.

Similarly, in order to study the effect under other modes of dominance, we have that the recursion for 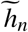 comes with *κ* = 2, therefore 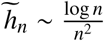. Then,

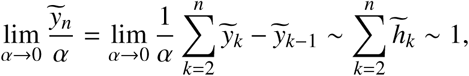

which gives the second and third assertion of Proposition 9.

## Acknowledgements

This research was supported by the DFG priority program SPP 1590, and in particular through grant Pf-672/8-1 to PP. We thank two anonymous referees for their careful reading and remarks, which helped to improve the manuscript.

